# Gonadal hormones and sex chromosome complement differentially contribute to ethanol intake, preference, and relapse-like behavior in Four Core Genotypes mice

**DOI:** 10.1101/2021.04.28.441845

**Authors:** Elizabeth A. Sneddon, Lindsay N. Rasizer, Natalie G. Cavalco, Asa H. Jaymes, Noah J. Ostlie, Brianna L. Minshall, Brianna M. Masters, Haley Hrncir, Arthur P. Arnold, Anna K. Radke

## Abstract

Alcohol use and high-risk alcohol drinking behaviors among women are rapidly rising. In rodent models, females typically consume more ethanol (EtOH) than males. Here, we used the Four Core Genotypes (FCG) mouse model to investigate the influence of gonadal hormones and sex chromosome complement on EtOH drinking behaviors. FCG mice were given access to escalating concentrations of EtOH in a two-bottle, 24-h continuous access drinking paradigm to assess consumption and preference. Relapse-like behavior was measured by assessing escalated intake following repeated cycles of deprivation and re-exposure. Twenty-four hour EtOH consumption was greater in mice with ovaries (*Sry*-), relative to those with testes, and in mice with the XX chromosome complement, relative to those with XY sex chromosomes. EtOH preference was higher in XX vs. XY mice. Escalated intake following repeated cycles of deprivation and re-exposure emerged only in XX mice (vs. XY). Mice with ovaries (*Sry*-FCG mice and C57Bl/6J females) were also found to consume more water than mice with testes. These results demonstrate that aspects of EtOH drinking behavior may be independently regulated by sex hormones and chromosomes and inform our understanding of the neurobiological mechanisms which contribute to EtOH dependence in male and female mice. Future investigation of the contribution of sex chromosomes to EtOH drinking behaviors is warranted.

## Introduction

Risky drinking behaviors and the development of Alcohol Use Disorder (AUD) is a prevalent health issue in the United States and worldwide ^1^. Recent research demonstrates that alcohol use and high-risk alcohol drinking behaviors among women are rapidly rising ^2^. Further, women may progress from initial alcohol experience to alcohol dependence more quickly than men ^3^. Paralleling the effects observed in humans, female rodents are known to be more vulnerable to a range of addictive behaviors ^4^. For example, female rodents consume more ethanol (EtOH) than males ^5–8^ and are more likely to consume EtOH despite the risk of punishment ^9–11^. The neurobiological mechanisms contributing to these behavioral differences are still relatively unknown.

One potential mediator of sex differences in alcohol drinking behaviors is gonadal hormones. Elevated levels of estrogens have been associated with higher levels of alcohol consumption in adolescent ^12^ and adult women ^13^. Recent work also suggests that rising progesterone levels may protect against alcohol intake in some women and that gonadal hormone levels may be associated with drinking alcohol to cope with negative emotional states ^14^. In rodents, elevated levels of EtOH consumption and EtOH reward in females ^15–17^ are at least partially dependent on ovarian hormones including estradiol, which promotes drinking ^18–23^ and EtOH reward ^24^.

Gonad type is influenced by the presence or absence of the *Sry* gene, which is located on the Y chromosome and is responsible for the development of testes in male mice and the secretion of testosterone ^25,26^. As such, gonad differentiation is the result of only one of a number of chromosomal differences between males and females. Indeed, the human Y chromosome encodes 27 proteins and the X chromosome encodes approximately 1500 proteins that could potentially contribute to sex differences in alcohol drinking behaviors and a large number of X-linked genes are important for brain development and function ^27–29^. Differentiating the influences of gonadal hormones and sex chromosomes on behavior is, however, difficult and there are few data on sex chromosome contributions to alcohol drinking.

Fortunately, the Four Core Genotypes (FCG) mouse model, in which the *Sry* gene is absent from the Y chromosome and inserted onto an autosome in the same mice, allows the influences of sex chromosomes and gonadal hormones on behavior to be assessed independently ^25,26^. Without the *Sry* transgene, XY or XX mice have female gonads (ovaries) regardless of sex chromosome complement, whereas XX or XY *Sry*+ mice have male gonads (testes) (**Fig. 1**). This results in four groups: XX/*Sry*-, XY/*Sry*-, XX/*Sry*+, and XY/*Sry*+. Using this mouse model, sex chromosomes have been found to influence aggression, parenting, nociception, social interaction, and habit formation ^25,30^. In regard to EtOH drinking, female gonads (*Sry*-) have been associated with increased consumption (paralleling the effects of gonadal hormones described above) while sex chromosomes influenced the development of habitual vs. goal-directed responding for EtOH ^31^.

**Figure 1.**
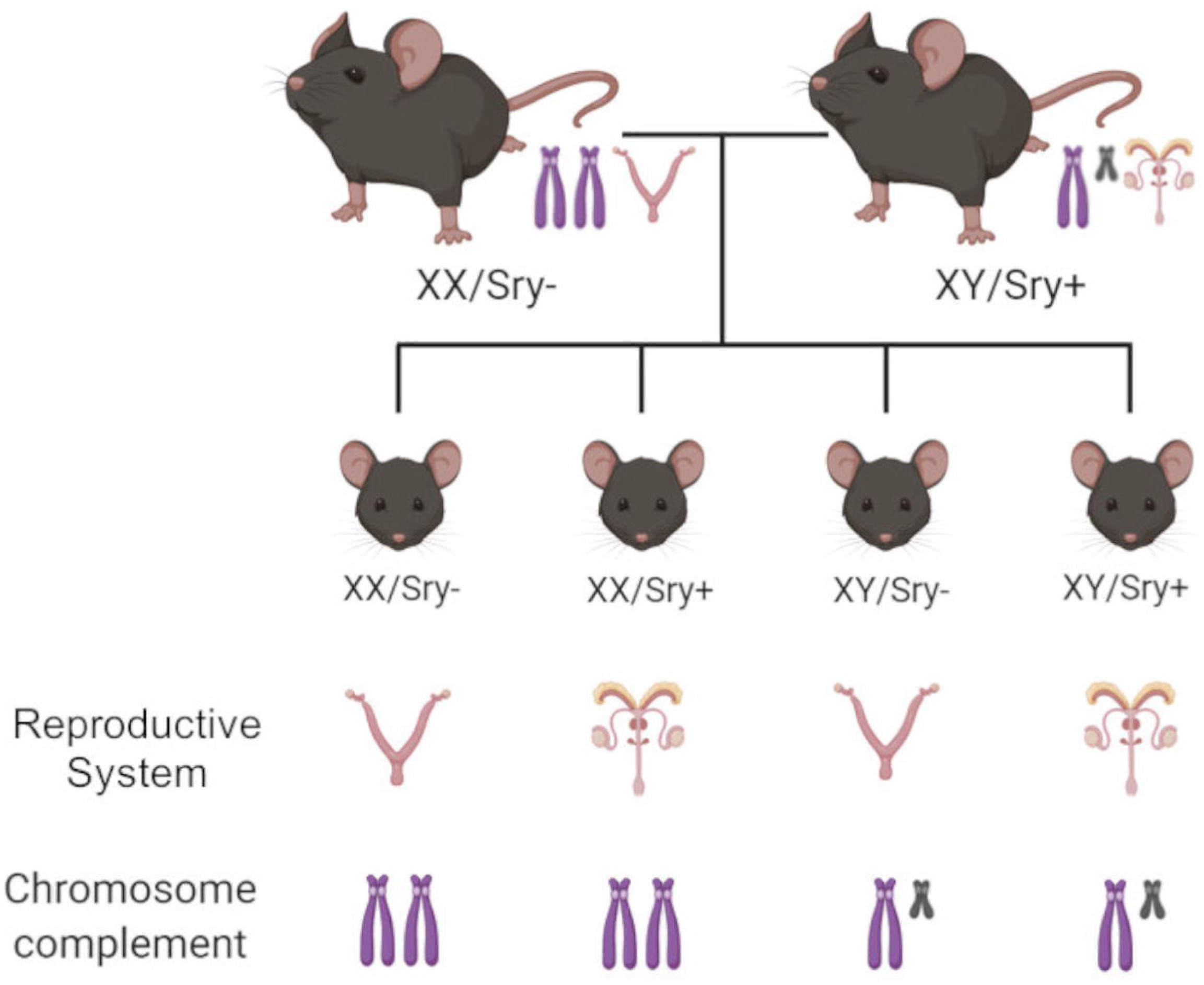
Breeding scheme of the Four Core Genotypes (FCG) mice. Breeding XX (wild-type) females with a XY/*Sry*+ mice results in four groups of offspring. XX/*Sry*- and XY/*Sry*-mice have ovaries and secrete female levels of hormones regardless of chromosome complement. XX/*Sry*+ and XY/*Sry*+ mice have testes and secrete male levels of hormones regardless of chromosome complement. Key: Purple chromosomes: mouse X chromosomes; gray chromosomes: mouse Y chromosomes; for *Sry*- mice the female mouse reproductive system is shown and for *Sry*+ mice the male mouse reproductive system is shown. The reproductive systems consist of gonadal (ovaries or testes) and non-gonadal tissues.

To explore the contributions of gonadal hormones and sex chromosomes to alcohol consumption, preference, and relapse-like behavior, we examined 24-h EtOH intake and the alcohol deprivation effect in FCG mice. Our results suggest an important role for both sex hormones and chromosomes in mediating female vulnerability to EtOH drinking behaviors.

## Methods

### Subjects

39 FCG mice (PND 60+) were generated from breeding pairs consisting of *Sry*+ XY male and C57BL/6J (wild type) XX female mice at the Laboratory of Animal Resources at Miami University ^26^. FCG breeding pairs used to generate experimental mice were obtained from UCLA. This breeding scheme results in four groups XX/*Sry*- (n = 10), XY/*Sry*- (n = 10), XX/*Sry*+ (n = 9), and XY/*Sry*+ (n = 10) (**Fig. 1**). Mice were tested in adulthood (XX/*Sry*-: 27.05 ± 1.85; XY/*Sry*-: 25.15 ± 2.46; XX/*Sry*+: 18.57 ± 1.63; XY/*Sry*+: 20.12 ± 1.84 weeks at the start of testing [all data mean ± SEM]). Prior to experimentation, FCG mice were group housed by gonad type (*Sry*- or *Sry*+). 52 C57BL/6J mice (male = 20, female = 32) were generated from breeding pairs purchased from the Jackson Laboratory (Bar Harbor, ME) and were grouped housed prior to experimentation or surgery.

One week before the first experimental session, mice were individually housed in standard shoe box udel polysufone rectangular mouse cages (18.4 cm × 29.2 cm × 12.7 cm) outfitted with 2-bottle cage tops. Mice were given standard care and had access to LabDiet 5001 standard chow and reverse-osmosis (RO) filtered water *ad libitum*. Mice were kept on a 12:12 hour light:dark cycle (lights on at 7 AM). All mice were cared for in accordance with the guidelines set by the National Institute of Health and all procedures were approved by the Institutional Animal Care and Use Committee (IACUC) at Miami University.

### 24-h home cage EtOH drinking and deprivation

Throughout testing, FCG mice had access to two bottles which contained reverse-osmosis (RO) drinking water and EtOH in RO water (v/v). Drinking bottles were made from 50 mL conical tubes fitted with ball-bearing sippers weighing approximately 80 g when filled and mice consumed about 2-4 g per session. All bottles were weighed every 24 h using a portable balance (Fisher Science Education, Model: SLF103, readability: 0.001 g). Every 48 h mice were weighed and solutions were changed out. EtOH concentrations increased over drinking sessions from 5%, 10%, 15%, to 20% (5 drinking sessions/ concentration) (**Fig. 2A**). After the last 20% EtOH session, mice underwent a 6-day EtOH deprivation period. After the deprivation period, mice were reintroduced to 20% EtOH for 24 h. This cycle of deprivation and re-exposure was then repeated for a total of 5 deprivation sessions (**Fig. 3A**). Bottles were alternated daily to equate side biases. Two “dummy” cages were outfitted with bottles to account for spillage and evaporation.

**Figure 2.**
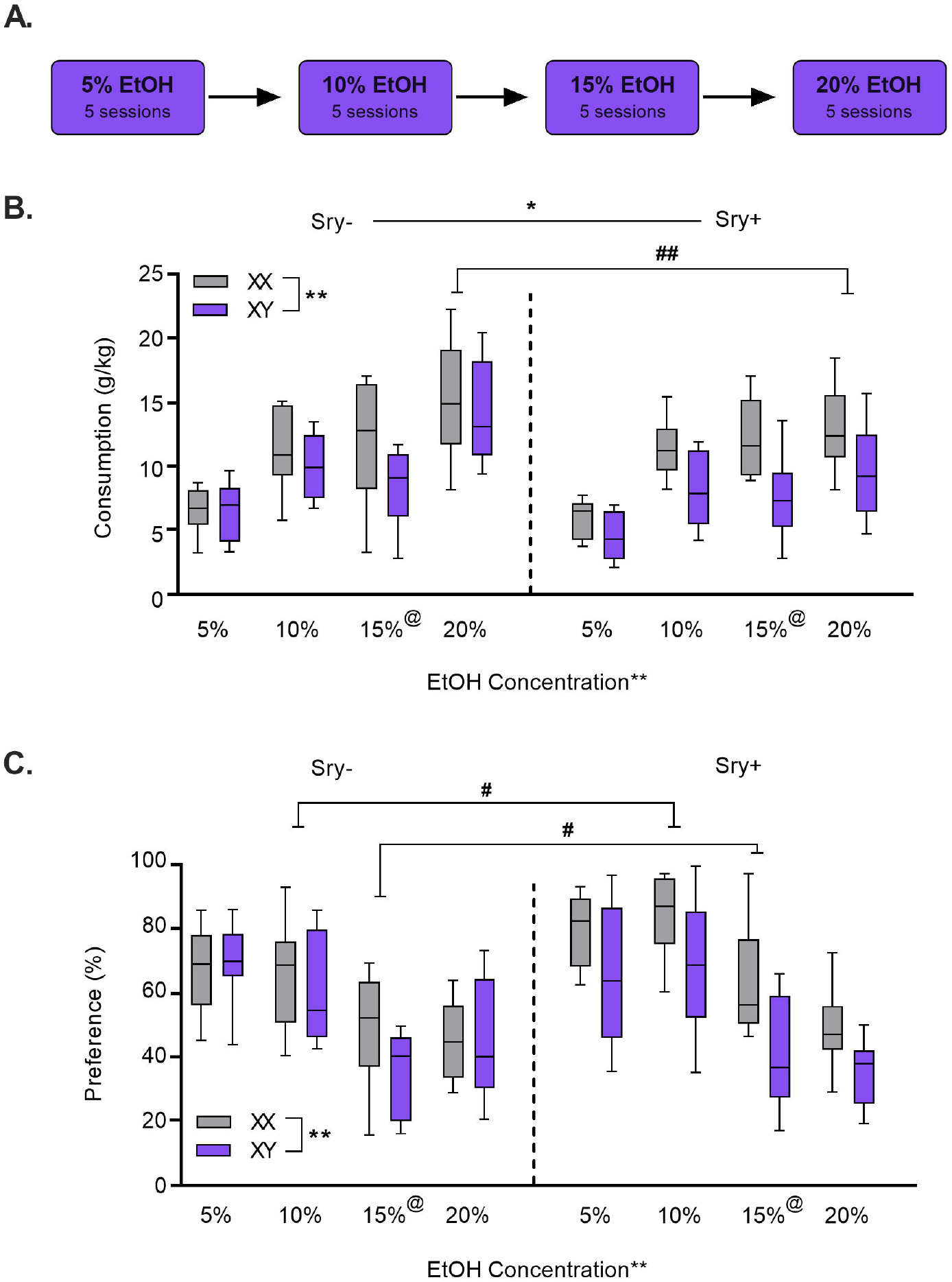
Differential effects of gonadal hormones and sex chromosomes on EtOH intake and preference. **A)** FCG mice drank EtOH, 5%, 10%, 15%, and 20% concentrations, for 24 h across 5 drinking sessions per concentration. **B)** *Sry*- (vs. *Sry*+) and XX (vs. XY) mice consumed greater amounts of EtOH. **C)** XX chromosomes were associated with heightened preference for EtOH vs. water. *p < 0.05, **p < 0.01 (main effects 3-Way ANOVA). ^#^p < 0.05, ^##^p < 0.01 main effect of *Sry*, and ^@^p < 0.01 main effect of chromosomes (for 2-Way ANOVA at that concentration).

**Figure 3.**
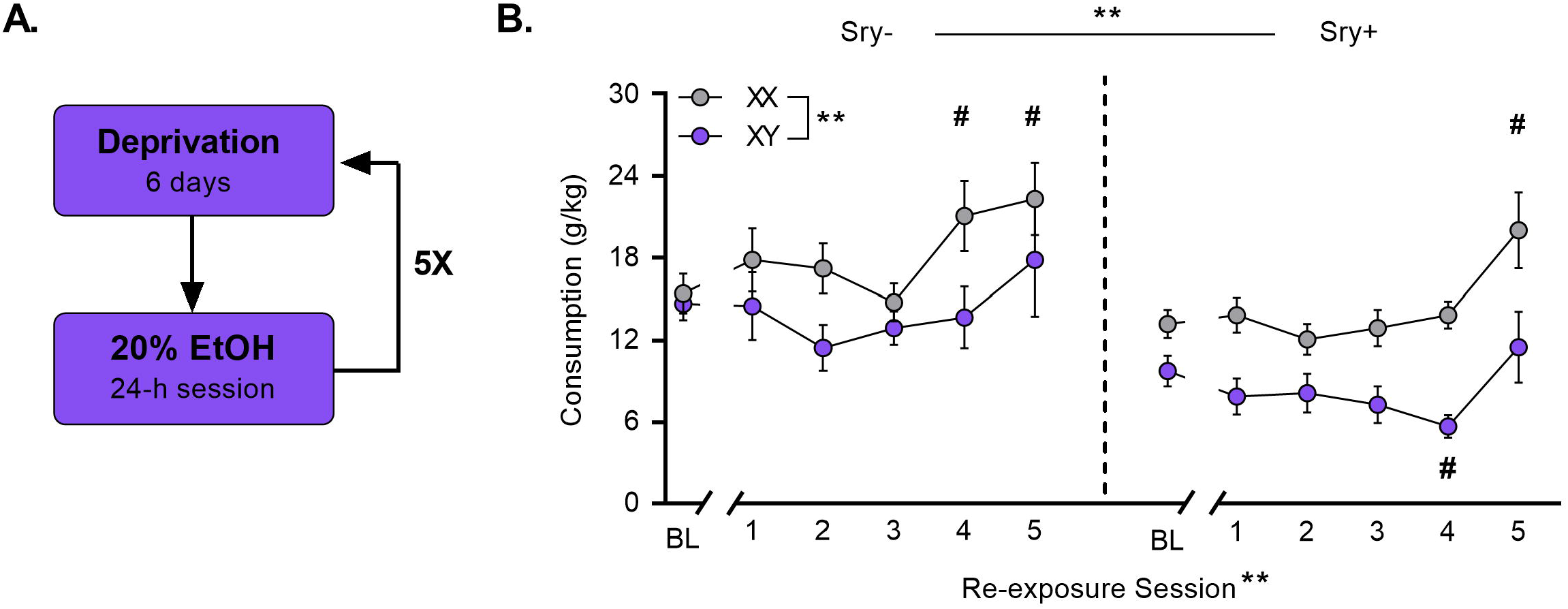
XX chromosomes promote the alcohol deprivation effect. **A)** Following the last 20% EtOH session, FCG mice underwent 6 sessions of deprivation and were then re-exposed to 20% EtOH for one 24-h session. This cycle of deprivation and re-exposure was repeated five times. **B)** XX/*Sry*- and XX/*Sry*+ mice escalated intake compared to drinking at baseline (= 5 sessions preceding deprivation; BL). # p < 0.05 vs. BL (Dunnett’s), **p < 0.01 (main effects 3-Way ANOVA).

### 24-h home cage water and sucrose drinking in FCG mice

At least two weeks following the last re-exposure session, FCG mice were presented with RO water for one 24-h session. The following day mice were presented with a bottle of 2.5% sucrose or RO water for a total of five 24-h drinking sessions (**Fig. 4A**).

**Figure 4.**
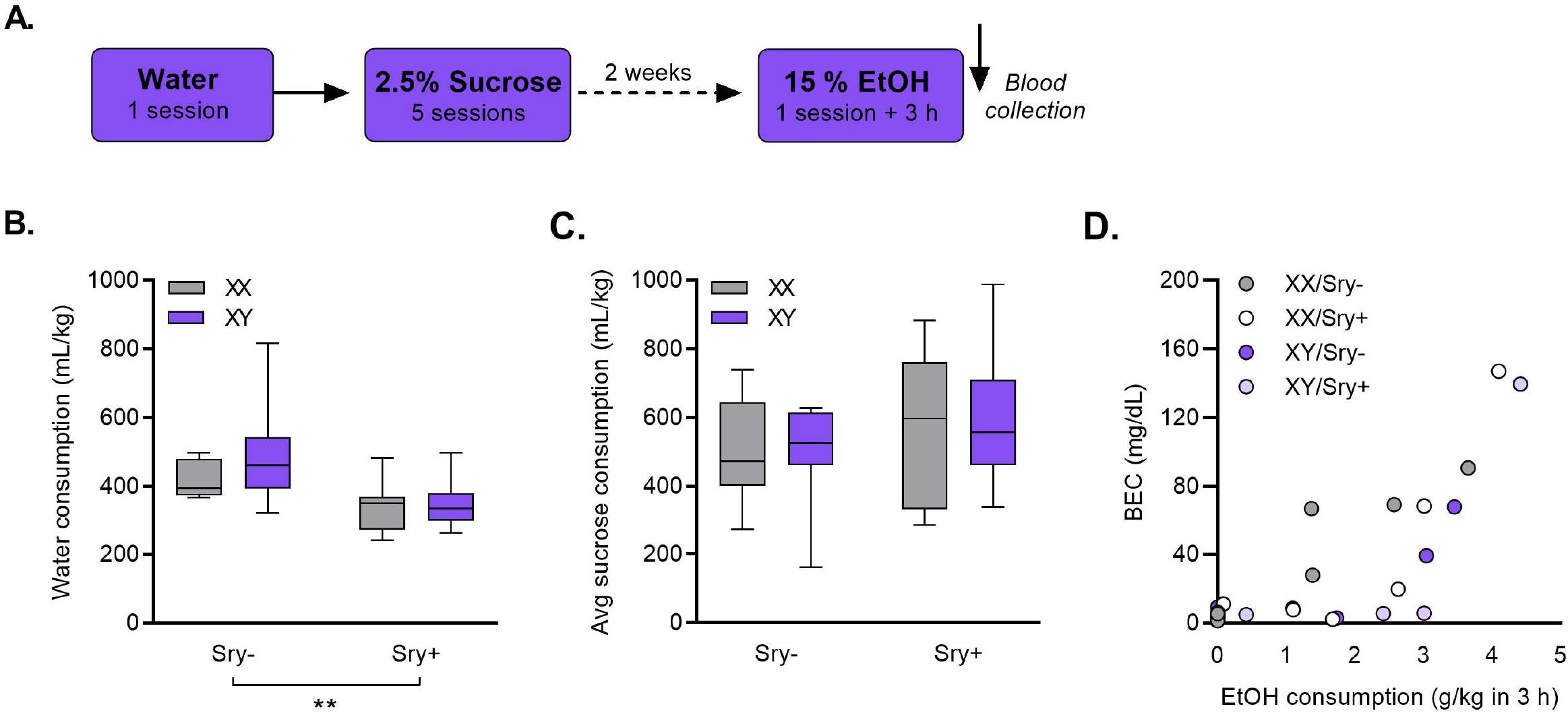
Female gonads (*Sry*-) promote water but not sucrose intake. **A)** FCG mice were given access to water for a 24-h session followed by five sessions with 2.5% sucrose. Two weeks later mice were reexposed to 15% EtOH for one 27-h session and blood samples were collected immediately after. **B)** *Sry*-mice consumed more water than *Sry*+ mice. ** p < 0.01 main effect of *Sry* (2-Way ANOVA). **C)** No differences were observed in sucrose consumption. **D**) 3-h consumption of 15% EtOH was significantly correlated with BEC measurements (p < 0.001).

### Blood EtOH concentration (BEC) analysis

At least two weeks following sucrose consumption (**Fig. 4A**), a subset of FCG mice (n = 28, XX/*Sry*- = 7, XY/*Sry*- = 7, XX/*Sry*+ = 7, XY/*Sry*+ = 7) were given access to 15% EtOH at the onset of the dark cycle (i.e., 7:00 PM) for a total of 27 h. Bottles were weighed 24 h into the session and at the end of the session. At the end of the session, blood samples were obtained by using the modified tail-clip technique ^32^. Mice were placed on a cage top and were allowed to explore while the researcher used a sterile razor blade to remove 1 mm from the tip of the tail. A heparinized capillary tube was used to collect 70 μL of blood. Blood was dispensed into a centrifuge tube and placed on ice. Immediately following collection, the blood was spun in an accuSpin Micro 17 microcentrifuge (Fisher Scientific) at 3000 rpm for 8 min. The serum was then pipetted out into a centrifuge tube and stored at −80 °C until testing. BECs were assessed using an AM1 alcohol analyzer (Analox Technologies).

### 24-h home cage water drinking in C57BL/6J mice

To follow up on effects observed in the FCG mice, water intake in C57BL/6J mice was assessed in two separate experiments. First, water intake in intact male and female mice was recorded over a 24-h session. Mice (n = 40, male = 20, female = 20) were individually housed at least 3 days prior to experimentation. One water bottle was presented to the mice *ab libitum* for one 24-h session. Bottles and mice were weighed at the onset and end of the session. Two “dummy” bottles were outfitted on cages to account for any evaporation or spillage.

The influence of circulating ovarian hormones on water intake was assessed in a separate experiment. Female C57BL/6J mice received either ovariectomy (OVX = 16) or sham (SHAM = 16) surgeries as described below. Water intake was assessed over a 24-h session as described above.

### Ovariectomy surgeries

Mice were placed under light anesthesia using isoflurane and were surgically prepped by removing the fur from the back. The incision sites were cleaned with three alternating swabs of betadine-soaked gauze and EtOH wipes. Next, a 1-cm incision was made through the skin along the abdominal cavity. The ovarian fat pat was identified under the dorsal muscle mass and a 0.5-cm incision was made through the dorsal muscle mass. The ovarian fat pad was pulled through the incision to expose the ovary. Hemostats were then used to clamp the uterine horn off underneath the ovary to prevent bleeding. A scalpel was used to remove the ovary. Following removal, the uterine horn and fat pad were replaced back into the body cavity. Sutures were made through the muscle and the skin to close the wound. The above steps were repeated for the next side. For sham surgeries, the above steps were the same but the ovary was not removed. After surgery, mice were placed on a heating pad for at least 30 minutes and were weighed before returning to the colony room.

Standard post-operative care was given for 3 – 5 days and 50 – 80 mg/kg/day of ibuprofen was available in drinking water. Experimentation occurred three weeks following surgery (after Satta et al., 2018).

### Data Analysis

Consumption was calculated as *(Initial Bottle Weight – Post Bottle Weight) – Average of Dummy Bottles*. EtOH consumption was expressed as grams of EtOH consumed per kilogram of body weight. Preference was calculated as ((*Volume of EtOH*)/ (*Volume of EtOH* + *Water Consumption*))*100. Consumption and preference for each concentration was computed by averaging across the five drinking sessions. Total water or EtOH consumption was calculated by summing consumption across all drinking sessions for each individual mouse then averaging for each group. Water and sucrose data were expressed as milliliters of water consumed per kilogram of body weight.

For EtOH consumption and preference data, a Three-Way ANOVA was used with *Sry* (gonad type) and sex chromosome complement as between-subjects factors and EtOH concentration as the within-subjects factor. Because *Sry* and chromosome complement were found to interact with EtOH concentration, but not each other, follow-up analyses using a Two-Way ANOVA were performed on each separate concentration using sex chromosome complement and *Sry* (gonad type) as between-subjects factors. In cases where the assumption of sphericity was violated (ε < 0.75), the Greenhouse-Geisser correction was applied. Dunnett’s test corrected for multiple comparisons was used to assess the alcohol deprivation effect, defined as an increase in consumption on re-exposure sessions vs. at baseline (= the five sessions immediately preceding the beginning of the first deprivation period). For water and sucrose consumption experiments in FCG mice, Two-Way ANOVA was used with *Sry* and chromosome complement as between-subject factors. In C57BL/6J mice, an unpaired t-test was used between groups. To correlate BECs with consumption, the Spearman’s rho correlation was used with 24-h intake and BEC level as factors.

Eta squared and Cohen’s d were calculated to report effect sizes as appropriate. All data are shown as mean ± SEM and the alpha level for all comparisons was set at *p* < 0.05. All analyses were conducted in GraphPad Prism v. 8.3 (La Jolla, CA). All images were created using GraphPad Prism, BioRender, and GNU Image Manipulation Program (GIMP) 2.10.

## Results

### Sex hormones and chromosomes influence EtOH consumption

Analysis of EtOH consumption across concentrations revealed influences of both *Sry* (gonad type) and sex chromosomes (**Fig. 2B**). *Sry*- mice consumed more EtOH than *Sry*+ mice and XX mice consumed more than XY mice. A Three-Way ANOVA identified main effects of the *Sry* gene (F_(1, 35)_ = 4.704, p = 0.037, η^2^ = 3.378%), chromosome complement (F_(1, 35)_ = 9.839, p = 0.004, η^2^ = 7.066%), and EtOH concentration (F_(2.439, 85.353)_ = 55.046, p < 0.0001, η^2^ = 36.086%). There were interactions with EtOH concentration for the *Sry* gene (F_(3, 105)_ = 3.156, p = 0.028, η^2^ = 2.071%) and chromosome complement (F_(3, 105)_ = 2.838, p = 0.042, η^2^ = 1.860%). The Three-Way interaction of *Sry* gene X sex chromosomes X EtOH concentration did not reach significance (F_(3, 105)_ = 0.237, p = 0.870, η^2^ = 0.156%). A follow-up Two-Way ANOVA found a main effect of chromosome complement (F_(1,35)_ = 12.645, p = 0.001, η^2^ = 26.367%) at the 15% EtOH concentration but no effect of *Sry* gene (F_(1,35)_ = 0.124, p = 0.727, η^2^ = 0.259%) or interaction (F_(1,35)_ = 0.177, p = 0.677, η^2^ = 0.368%). A follow-up Two-Way ANOVA discovered a main effect of *Sry* gene (F_(1,35)_ = 8.222, p = 0.007, η^2^ = 17.235%) at the 20% EtOH concentration but no effect of chromosome complement (F_(1,35)_ = 2.928, p = 0.096, η^2^ = 6.138%) or interaction (F_(1,35)_ = 1.213, p = 0.278, η^2^ = 2.542%).

Additional analyses on body weights across EtOH concentrations revealed that *Sry*+ mice weighed more than *Sry*-mice at the 10%, 15%, and 20% concentrations (**Table 1**). A Three-Way ANOVA identified a main effect of the *Sry* gene (F_(1,35)_ = 10.706, p = 0.002, η^2^ = 14.299%) and an interaction between EtOH concentration X *Sry* gene (F_(3,105)_ = 3.512, p = 0.018, η^2^ = 3.086%). The Three-Way interaction of *Sry* gene X sex chromosomes X EtOH concentration did not reach significance (F_(3,105)_ = 0.249, p = 0.862, η^2^ = 0.219%). A follow-up Two-Way ANOVA showed a significant main effect of *Sry* gene (F_(1,35)_ = 14.002, p < 0.001, η^2^ = 27.096%) at the 10% EtOH concentration but no effect of chromosome complement (F_(1,35)_ = 2.068, p = 0.159, η^2^ = 4.002%) or interaction (F_(1,35)_ = 0.616, p = 0.427, η^2^ = 1.251%).

**Table 1:** Average body weights of Four Core Genotypes mice. Mice with testes weighed more than mice with ovaries. *Sry*+ weighed more than *Sry*-mice at the 10%, 15%, and 20% concentrations. ^**a**^ main effect of *Sry* gene, p < 0.01 (2-Way ANOVA). Data are averages ± standard error of the mean (SEM).

| Group | 5% EtOH (mL/ kg) | 10% EtOH (mL/ kg)<sup>a</sup> | 15% EtOH (mL/ kg)<sup>a</sup> | 20% EtOH (mL/ kg)<sup>a</sup> |
| --- | --- | --- | --- | --- |
| XX/Sry- | 24.25 ± 0.52 | 22.68 ± 0.35 | 23.95 ± 0.39 | 23.94 ± 0.49 |
| XY/Sry- | 23.40 ± 0.54 | 22.62 ± 0.52 | 23.07 ± 0.58 | 23.11 ± 0.49 |
| XX/Sry+ | 23.85 ± 0.32 | 24.96 ± 0.43 | 25.08 ± 0.43 | 25.22 ± 0.43 |
| XY/Sry+ | 24.32 ± 0.66 | 24.68 ± 0.48 | 24.76 ± 0.52 | 25.02 ± 0.47 |

### Sex hormones and chromosomes influence preference for EtOH vs. water

Analysis of EtOH preference demonstrated an influence of sex chromosomes and hormones (**Fig. 2C**). Preference was higher in XX (vs. XY mice) and, at the 10% and 15% concentrations, in *Sry*+ mice (vs. *Sry*-). A Three-Way ANOVA revealed a main effect of chromosome complement (F_(1, 35)_ = 8.130, p = 0.007, η^2^ = 5.784%) and EtOH concentration (F_(2.248, 78.673)_ = 47.188, p < 0.0001, η^2^ = 35.168%), but no interaction. The main effect of *Sry* gene was not significant (F_(1, 35)_ = 3.136, p = 0.085, η^2^ = 2.232%) but there was a significant *Sry* gene X EtOH concentration interaction (F_(3, 105)_ = 3.169, p = 0.027, η^2^ = 2.362%). The interactions between *Sry* gene and sex chromosomes (F_(1,35)_ = 3.555, p = 0.068, η^2^ = 2.529%) and *Sry* gene, sex chromosomes, and EtOH concentrations (F_(3,105)_ = 0.117, p = 0.950, η^2^ = 0.087%) did not reach significance. A follow-up Two-Way ANOVA discovered a main effect of *Sry* gene (F_(1,35)_ = 6.390, p = 0.0161, η^2^ = 14.129%) at the 10% EtOH concentration but no effect of chromosome complement (F_(1,35)_ = 3.016, p = 0.091, η^2^ = 6.669%) or interaction (F_(1,35)_ = 1.294, p = 0.263, η^2^ = 2.861%). A follow-up Two-Way ANOVA found a main effect of chromosome complement (F_(1,35)_ = 13.964, p = 0.001, η^2^ = 25.938%) and *Sry* gene (F_(1,35)_ = 4.403, p = 0.043, η^2^ = 8.179%) at the 15% EtOH concentration but no interaction (F_(1,35)_ = 1.191, p = 0.2827, η^2^ = 2.211%).

Higher EtOH consumption in *Sry*-mice without corresponding increases in EtOH preference suggested that gonad type also influences water consumption (**Table 2**). A Three-Way ANOVA of water consumption during the 24-h EtOH drinking paradigm in *Sry*- and *Sry*+ mice demonstrated that *Sry*-mice consumed more water than *Sry*+ mice, as evidenced by a main effect of *Sry* gene (F_(1, 35)_ = 13.338, p < 0.001, η^2^ = 13.953%) and of EtOH concentration (F_(2.171,75.991)_ = 30.486, p < 0.0001, η^2^ = 19.963%). The Three-Way interaction of *Sry* gene X sex chromosomes X EtOH concentration did not reach significance (F_(3, 105)_ = 0.258, p = 0.855, η^2^ = 0.169%). A follow-up Two-Way ANOVA discovered a main effect of *Sry* gene (F_(1,35)_ = 6.351, p = 0.0164, η^2^ = 14.446%) at the 5% EtOH concentration but no effect of chromosome complement (F_(1,35)_ = 0.161, p = 0.501, η^2^ = 1.050%) or interaction (F_(1,35)_ = 2.494, p = 0.123, η^2^ = 5.673%). A follow-up Two-Way ANOVA revealed a main effect of *Sry* gene (F_(1,35)_ = 11.444, p = 0.002, η^2^ = 24.101%) at the 10% EtOH concentration but no effect of chromosome complement (F_(1,35)_ = 1.052, p = 0.312, η^2^ = 2.216%) or interaction (F_(1,35)_ = 0.275, p = 0.604, η^2^ = 0.578%). A follow-up Two-Way ANOVA found a main effect of chromosome complement (F_(1,35)_ = 8.924, p = 0.005, η^2^ = 0.365%) and *Sry* gene (F_(1,35)_ = 15.117, p < 0.0001, η^2^ = 25.836%) at the 15% EtOH concentration but no interaction (F_(1,35)_ = 0.213, p = 0.647, η^2^ = 0.365%).

**Table 2.** Water consumption in Four Core Genotypes mice. Gonad type predicted water consumption. *Sry*-mice consumed more water when 5% or 10% EtOH was present. *Sry*- (vs. *Sry*+) and XY (vs. XX) mice consumed more water when 15% EtOH was present. ^**a**^ main effect of *Sry* gene, p < 0.05 (2-Way ANOVA). ^**b**^ main effect of chromosomes, p = 0.05 (2-Way ANOVA) Data are averages ± standard error of the mean (SEM).

| Group | 5% EtOH (mL/ kg)<sup>a</sup> | 10% EtOH (mL/ kg)<sup>a</sup> | 15% EtOH (mL/ kg)<sup>ab</sup> | 20% EtOH (mL/ kg) |
| --- | --- | --- | --- | --- |
| XX/Sry- | 88.33 ± 13.93 | 88.65 ± 15.66 | 120.02 ± 11.21 | 129.78 ± 14.40 |
| XY/Sry- | 77.95 ± 8.45 | 95.82 ± 17.06 | 153.91 ± 19.45 | 139.10 ± 25.69 |
| XX/Sry+ | 41.02 ± 6.07 | 32.79 ± 10.70 | 61.65 ± 9.93 | 90.84 ± 8.06 |
| XY/Sry+ | 67.09 ± 14.46 | 54.95 ± 12.07 | 107.94 ± 9.92 | 118.12 ± 4.49 |

### Sex chromosomes influence the alcohol deprivation effect

Mice were re-exposed to 20% EtOH following 6-day cycles of deprivation for 5 total re-exposures (**Fig. 3A**). Previous research suggests that this regimen promotes increases in intake known as the alcohol deprivation effect ^33^. Analysis of EtOH consumption at baseline (= the five sessions immediately preceding the beginning of the first deprivation period) and on the five deprivation sessions demonstrated that sex chromosomes influenced the magnitude of the alcohol deprivation effect. A Three-Way ANOVA revealed main effects of the *Sry* gene (F_(1, 35)_ = 12.075, p = 0.001, η^2^ = 10.953%), sex chromosome complement (F_(1, 35)_ = 13.202, p = 0.0009, η^2^ = 11.975%), and re-exposure session (F_(3.119, 109.174)_ = 7.947, p < 0.0001, η^2^ = 7.618%). No interactions were observed between re-exposure session X *Sry* gene (F_(5,175)_ = 0.969, p = 0.438, η^2^ = 0.929%), re-exposure session X chromosome complement (F_(5,175)_ = 1.719, p = 0.133, η^2^ = 0.493%), *Sry* gene X chromosome complement (F_(1,35)_ = 0.544, p = 0.466, η^2^ = 0.493%), or re-exposure session X *Sry* gene X chromosome complement (F_(5,175)_ = 0.521, p = 0.760, η^2^ = 0.466%).

Follow up Dunnett’s tests (control condition = baseline) showed that mice in the XX/*Sry*-group consumed more EtOH on re-exposure sessions 4 (p = 0.039) and 5 (p = 0.047) and mice in the XX/*Sry*+ group consumed more EtOH on re-exposure session 5 (p = 0.039) compared to baseline (**Fig. 3B**). Mice in the XY/*Sry*+ group consumed less EtOH compared to baseline on re-exposure session 4 (p=0.032).

### Sex hormones influence water but not sucrose consumption

To determine whether the effects of the *Sry* gene on water consumption could be replicated without concurrent access to EtOH, water consumption in the home cage during a 24-h period was measured in all FCG mice. A Two-Way ANOVA revealed that *Sry*-mice drank more water vs. *Sry*+ mice, as evidenced by a main effect of *Sry* (F_(1, 35)_ = 13.339, p < 0.001, η^2^ = 26.099%) (**Fig. 4B**). The 95% CI was (−99.857, 21.983) between chromosomes and (48.677, 170.517) between gonad type.

A final control experiment was conducted to assess whether the observed differences in EtOH consumption and preference were specific to EtOH reward. FCG mice drank a 2.5% sucrose solution in a two-bottle choice paradigm for five 24-h sessions. Two-Way ANOVAs identified no group differences in average consumption (**Fig. 4C**) or preference for sucrose.

### Blood EtOH concentrations did not differ between genotypes

Two weeks following sucrose consumption, a subset of FCG mice were re-exposed to 15% EtOH for 27 h and blood samples were collected at the end of the session. A Two-Way ANOVA revealed no significant differences in consumption between groups at the 24 h or 3 h timepoints. BECs were significantly correlated with 3-h consumption (*r_s_* = 0.581, p = 0.001) (**Fig. 4D**). A Two-Way ANOVA also identified no significant differences in BECs between groups (XX/*Sry*-: 38.31 ± 13.91, XX/*Sry*+: 37.18 ± 20.28, XY/*Sry*-: 18.93 ± 9.54, XY/*Sry*+: 24.27 ± 19.24 [all data mean mg/dl ± SEM]).

### Female mice consume more water than male mice

In one 24-h session, female C57BL/6J consumed more water than their male counterparts. An unpaired t-test identified a difference between male and female consumption (t_(38)_ = 2.933, p = 0.005, d = 0.849) (**Fig. 5A**). The 95% CI was (19.133, 104.381) between factors.

**Figure 5.**
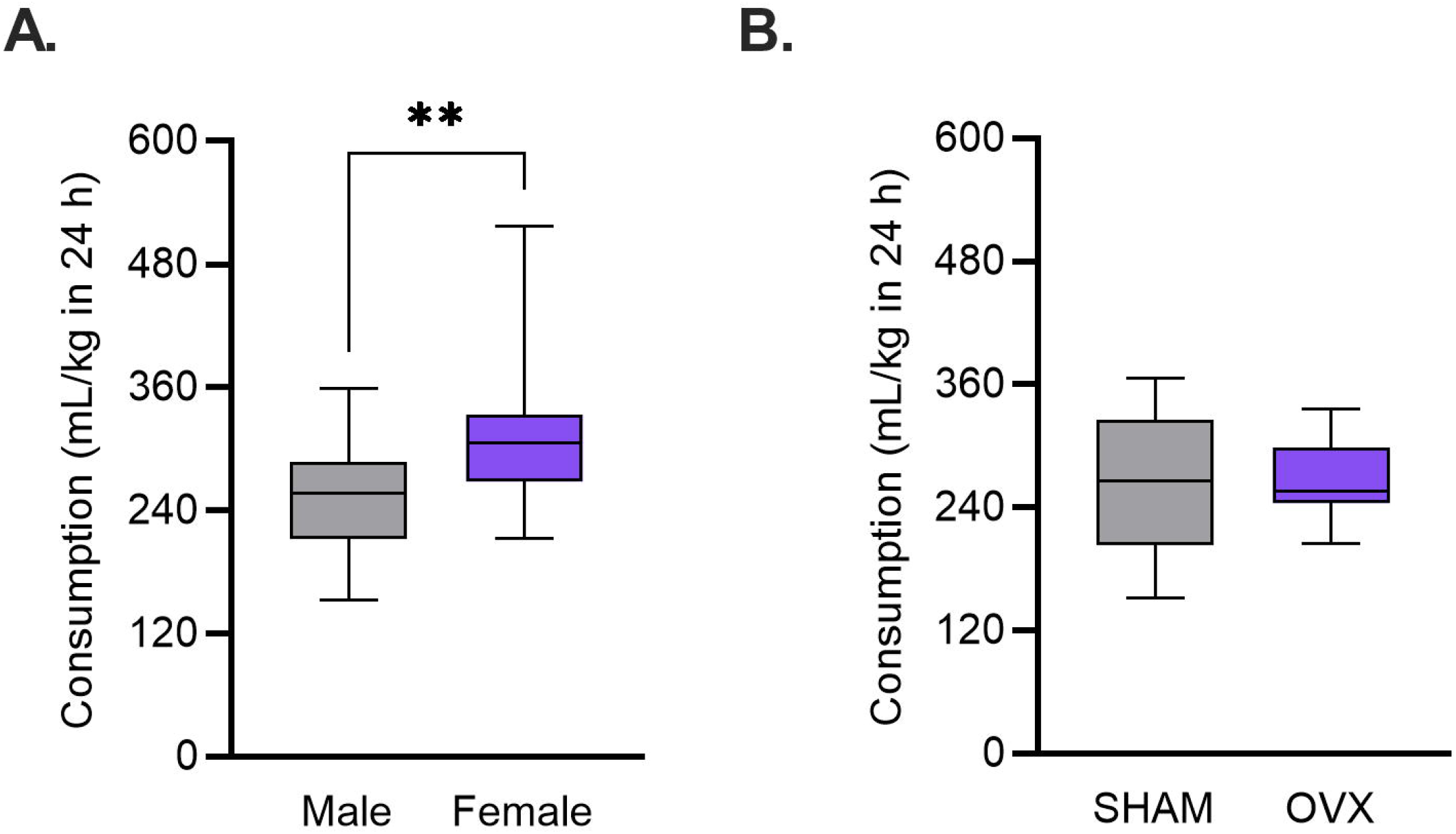
Circulating ovarian hormones do not influence water intake. C57BL/6J mice were given access to water for a 24-hour session. **A)** Female mice consumed more water than male mice. ** p < 0.01 (Unpaired t-test). **B)** Ovariectomized (OVX) mice consumed similar amounts of water compared to controls (SHAM).

### Water consumption is not dependent on circulating ovarian hormones

In one 24-h session, water consumption in ovariectomized C57BL/6J mice was unchanged compared to controls. An unpaired t-test found no difference between OVX and SHAM water consumption (t_(30)_ = 0.213, p = 0.833, d = 0.077) (**Fig. 5B**). The 95% CI was (−35.63, 43.95) between factors.

## Discussion

Using the FCG mouse model in a 24-h continuous access drinking paradigm, we explored how sex hormones and sex chromosomes contribute to female vulnerability to EtOH consumption, preference, and relapse-like behavior and uncovered novel influences of sex chromosomes. Twenty-four-hour EtOH consumption was greater in mice with female gonads (*Sry*-) and in mice with the XX chromosome complement. EtOH preference was higher in XX vs. XY mice. Escalated intake following repeated cycles of deprivation and re-exposure emerged only in XX mice (vs. XY). Follow-up control studies demonstrated that *Sry*-mice consumed more water than their *Sry*+ counterparts did and that drinking resulted in similar blood EtOH levels among the genotypes. These results concur with prior findings concerning the role of ovarian hormones in female vulnerability to EtOH drinking while revealing an underappreciated role for sex chromosomes in these behaviors.

Because the presence or absence of *Sry* determines the development of ovaries (*Sry*-) vs. testes (*Sry*+), behavioral differences between the *Sry* genotypes are primarily driven by levels of gonadal hormones. It is well established that ovarian hormones such as estradiol promote consumption and seeking of alcohol in rats and mice ^15,18–21,23^. Our observation of higher intake in *Sry*-mice similarly suggests that female gonads promote EtOH consumption. This finding also replicates an earlier study in FCG mice that reported higher 30-min EtOH consumption in *Sry*-mice ^31^. In the current study, the effect of *Sry* on consumption was most pronounced at the highest concentration of EtOH (20%), which supports a previous study from our lab that found higher consumption in female vs. male C57BL/6J mice only at concentrations above 10% ^11^. Thus, the current results provide further evidence that, at least in mice, female vulnerability to EtOH consumption is concentration dependent.

Interestingly, the increased EtOH consumption observed in *Sry*-mice was not seen for EtOH preference. In fact, at some concentrations, preference was higher in *Sry*+ mice (vs. *Sry*-). Some prior studies have seen increased preference in female rodents ^15^ while others have not ^8,10^. We have observed a similar pattern of greater EtOH consumption but not preference in intact female vs. male C57BL/6J mice using a drinking in the dark paradigm ^8^ and similar findings have been seen in rats ^34,35^. In the current study, the absence of an increase in preference in females was driven by elevations in water intake in *Sry*-mice during EtOH drinking sessions, relative to *Sry*+ mice. Higher water intake was also observed in a separate, EtOH-free session in *Sry*- vs. *Sry*+ mice and in female vs. male C57BL/6J mice. Although estrogens have been shown to influence water consumption in rodents ^36–38^, we did not find an influence of circulating ovarian hormones on water consumption in OVX vs. SHAM female C57BL/6J mice. The influence of gonad type on water intake is an important consideration for studies of EtOH drinking behaviors, since variations in water intake could influence blood EtOH levels. It also raises the possibility that mice with female gonads may simply drink more fluids. The observation of equal sucrose consumption across genotypes could argue against this conclusion, although we cannot rule out that consumption levels in that experiment reached a ceiling and obscured any effects.

In addition to gonadal influences on EtOH consumption, we report a novel role for sex chromosomes in influencing both consumption and preference. In one other study of 30-min consumption of 10% EtOH in FCG mice, sex chromosome complement did not influence consumption ^31^. In the current study, differences between XX and XY mice were greatest at the 15% concentration. Thus, the effects of chromosomes may be dependent on the drinking paradigm used or the concentration of EtOH presented. Importantly, sex chromosomes did not influence water or sucrose intake, suggesting that the observed effects are specific to EtOH. When considered alongside our findings regarding *Sry* influences on consumption but not preference, these results suggest that sex chromosome differences between males and females play a critical, and perhaps primary, role in driving sex differences in some EtOH drinking behaviors.

Another major finding of this work is that sex chromosomes and hormones influenced relapse-like behavior. As in the 24-h continuous portion of the experiment, EtOH consumption during re-exposure sessions was higher for mice with XX (vs. XY) chromosomes and *Sry*- (vs. *Sry*+) mice. Furthermore, escalations in drinking following deprivation (assessed by comparing consumption on re-exposure sessions to baseline consumption of 20% EtOH) were only observed in mice with XX chromosomes. The observation of the alcohol deprivation effect on re-exposure session 4 in XX/*Sry*-but not XX/*Sry*+ mice further suggests that the presence of female gonads facilitates the effects of deprivation on drinking behavior. One important consideration is that the alcohol deprivation effect was completely absent in XY/*Sry*+ mice, which are most similar to the male mice first used to establish this paradigm in the C57BL/6J line ^33^. Interestingly, delayed emergence of escalated drinking (during re-exposure session 6-7) was observed in a study by Melendez and colleagues (2006) when variants of the paradigm were used, including when a higher concentration of EtOH was available. It is possible that a similar delay occurred here in the XY/*Sry*+ mice because we used 20% EtOH, as opposed to 15%. Thus, while we cannot be certain, we think it likely that escalated drinking emerges more quickly (i.e., following fewer deprivation cycles) in vulnerable animals.

The alcohol deprivation effect has been demonstrated previously in male rats and mice ^33,39,40^ and has also been observed in female rats and WSC-1 mice following a single two-week deprivation ^41,42^. In the latter study, escalations in drinking following deprivation were similar in male and female mice ^42^. As such, our results are the first demonstration of greater female susceptibility in this model of relapse-like behavior. One limitation of our study is that we did not assess the deprivation effect by comparing consumption to non-deprived animals, as has been done previously ^33^. Instead, we defined the deprivation effect as an increase in drinking compared to baseline. It is also important to note that prior EtOH exposure may have influenced the results of subsequent control experiments assessing sucrose, water, and EtOH consumption. Indeed, gonad and chromosome effects on EtOH drinking were no longer apparent in a final control session performed in a subset of mice for the purpose of measuring blood EtOH levels.

Another important limitation of this study is that the estrous cycle was not assessed in *Sry*-mice. Estrous phase is typically unrelated to EtOH consumption levels in freely cycling rodents ^9,18,23,43,44^ supporting other work demonstrating that gonadal hormone effects on EtOH consumption in females occur during development (i.e., are organizational rather than activational) ^15,16,45,46^. Further, because we averaged intake and preference for each concentration of EtOH over 5 sessions, there are unlikely to be influences of estrous on the 24-h, continuous drinking results. We cannot, however, exclude the possibility that estrous cycle phase influenced results from the re-exposure sessions. Complicating the issue is the fact that extended EtOH exposure can alter the estrous cycle ^47–49^ and XY mice with ovaries stop cycling earlier in life than XX mice ^25^. A related consideration is that we do not have data on circulating hormone levels in the FCG mice. However, FCG mice with XX chromosomes are fully feminized and XY mice fully masculinized on a number of traits ^26^, and gonadal hormones levels have consistently been found to be similar in XX and XY FCG mice with the same type of gonads ^50,51^. Future studies in gonadectomized FCG mice will nevertheless be needed to fully resolve this issue as well as to confirm that the effects of gonadal hormones are organizational in nature.

A sex difference driven by sex chromosome complement may be due to genes on the Y chromosome, an extra dose of genes on the X chromosome in females, or paternal imprinting of X-linked genes ^52,53^. The mechanisms through which sex chromosomes influence EtOH drinking behaviors remain to be determined. Prior studies in the FCG mice have uncovered a role for sex chromosomes in the development of habit formation^331,54^, locomotor activation to cocaine ^55^, and in the density of tyrosine hydroxylase-expressing neurons in the midbrain ^52,56^ (though note that no effect on sucrose consumption was observed in the current study). It is thus possible that genes on the X and/or Y chromosomes produce sex differences in EtOH consumption, preference, relapse-like behavior, and/or habit formation via influences on the development of brain reward systems.

In sum, the current results confirm an involvement of gonadal hormones and highlight a currently underappreciated contribution of sex chromosomes in EtOH drinking behaviors. In addition to continued exploration of the mechanisms by which sex hormones facilitate EtOH drinking, it is critical that future studies begin to explore sex-specific genetic influences on these behaviors.

## Acknowledgments

This study was supported by NIH grants R15 AA027915 (AKR) and F99 NS118727 (EAS) and R01 HD076125 (APA) and by the College of Arts and Sciences, the Office of Research for Undergraduates, and the Department of Psychology’s Broadening Undergraduate Research Participation in Behavioral Neuroscience program at Miami University. The authors would like to thank Kristen M. Schuh for assistance with behavioral experiments and Catherine Wasylyshyn for assistance with surgeries. The authors declare no conflicting interests in this work.

## Author Contributions

AKR, EAS, LNR, NGC, ASJ, and NJO designed the experiments. EAS, LNR, NGC, ASJ, and NJO conducted the experiments. EAS and BLM conducted the surgeries. HH and APA provided the FCG breeding pairs. AKR, EAS, LNR, NGC, ASJ, and NJO analyzed data. AKR, EAS, and APA interpreted the findings. EAS drafted the manuscript. All authors edited, contributed to, and approved the manuscript.

## References

1. Alcohol Facts and Statistics. National Institute on Alcohol Abuse and Alcoholism. Published 10/2020. Accessed December 3, 2020. https://www.niaaa.nih.gov/publications/brochures-and-fact-sheets/alcohol-facts-and-statistics

2. Grant BF, Chou SP, Saha TD, et al. Prevalence of 12-month alcohol use, high-risk drinking, and DSM-IV alcohol use disorder in the United States, 2001-2002 to 2012-2013: Results from the National Epidemiologic Survey on Alcohol and Related Conditions. JAMA Psychiatry. 2017;74(9):911–923.

3. Anglin MD, Hser YI, McGlothlin WH. Sex differences in addict careers. 2. Becoming addicted. Am J Drug Alcohol Abuse. 1987;13(1-2):59–71.

4. Anker JJ, Carroll ME. Females are more vulnerable to drug abuse than males: evidence from preclinical studies and the role of ovarian hormones. Curr Top Behav Neurosci. 2011;8:73–96.

5. Lancaster FE, Spiegel KS. Sex differences in pattern of drinking. Alcohol. 1992;9(5):415–420.

6. Juárez J, Barrios de Tomasi E. Sex differences in alcohol drinking patterns during forced and voluntary consumption in rats. Alcohol. 1999;19(1): 15–22.

7. Middaugh LD, Kelley BM, Bandy ALE, McGroarty KK. Ethanol consumption by C57BL/6 mice: Influence of gender and procedural variables. Alcohol. 1999;17(3): 175–183.

8. Sneddon EA, White RD, Radke AK. Sex Differences in Binge-Like and Aversion-Resistant Alcohol Drinking in C57 BL/6J Mice. Alcohol Clin Exp Res. 2019;43(2):243–249.

9. Fulenwider HD, Nennig SE, Price ME, Hafeez H, Schank JR. Sex Differences in Aversion-Resistant Ethanol Intake in Mice. Alcohol Alcohol. 2019;54(4):345–352.

10. Radke AK, Held IT, Sneddon EA, Riddle CA, Quinn JJ. Additive influences of acute early life stress and sex on vulnerability for aversion-resistant alcohol drinking. Addict Biol. 2019;14(February): 1–10.

11. Sneddon EA, Ramsey OR, Thomas A, Radke AK. Increased Responding for Alcohol and Resistance to Aversion in Female Mice. Alcohol Clin Exp Res. Published online May 29, 2020. doi:10.1111/acer.14384

12. Martin CA, Mainous AG 3rd, Curry T, Martin D. Alcohol use in adolescent females: correlates with estradiol and testosterone. Am J Addict. 1999;8(1):9–14.

13. Muti P, Trevisan M, Micheli A, Krogh V, Bolelli G. Alcohol consumption and total estradiol in premenopausal women. Cancer Epidemiol. Published online 1998. https://cebp.aacrjournals.org/content/7/3/189.short

14. Holzhauer CG, Wemm SE, Wulfert E, Cao ZT. Fluctuations in progesterone moderate the relationship between daily mood and alcohol use in young adult women. Addict Behav. 2020;101:106146.

15. Almeida OF, Shoaib M, Deicke J, Fischer D, Darwish MH, Patchev VK. Gender differences in ethanol preference and ingestion in rats. The role of the gonadal steroid environment. J Clin Invest. 1998;101(12):2677–2685.

16. Cailhol S, Mormède P. Sex and strain differences in ethanol drinking: Effects of gonadectomy. Alcohol Clin Exp Res. 2001;25(4):594–599.

17. Torres OV, Walker EM, Beas BS, O’Dell LE. Female Rats Display Enhanced Rewarding Effects of Ethanol That Are Hormone Dependent. Alcohol Clin Exp Res. 2014;38(1): 108–115.

18. Ford MM, Eldridge JC, Samson HH. Ethanol consumption in the female Long–Evans rat. Alcohol. 2002;26(2):103–113.

19. Reid LD, Marinelli PW, Bennett SM, et al. One injection of estradiol valerate induces dramatic changes in rats’ intake of alcoholic beverages. Pharmacol Biochem Behav. 2002;72(3):601–616.

20. Reid ML, Hubbell CL, Reid LD. A pharmacological dose of estradiol can enhance appetites for alcoholic beverages. Pharmacol Biochem Behav. 2003;74(2):381–388.

21. Ford MM, Eldridge JC, Samson HH. Determination of an estradiol dose-response relationship in the modulation of ethanol intake. Alcohol Clin Exp Res. 2004;28(1):20–28.

22. Rajasingh J, Bord E, Qin G, et al. Enhanced voluntary alcohol consumption after estrogen supplementation negates estrogen-mediated vascular repair in ovariectomized mice. Endocrinology. 2007;148(8):3618–3624.

23. Satta R, Hilderbrand ER, Lasek AW. Ovarian Hormones Contribute to High Levels of Binge-Like Drinking by Female Mice. Alcohol Clin Exp Res. 2018;42(2):286–294.

24. Hilderbrand ER, Lasek AW. Estradiol enhances ethanol reward in female mice through activation of ERα and ERβ. Horm Behav. 2018;98(December 2017):159–164.

25. Arnold AP, Chen X. What does the “four core genotypes” mouse model tell us about sex differences in the brain and other tissues? Front Neuroendocrinol. 2009;30(1): 1–9.

26. De Vries GJ, Rissman EF, Simerly RB, et al. A model system for study of sex chromosome effects on sexually dimorphic neural and behavioral traits. J Neurosci. 2002;22(20):9005–9014.

27. Zechner U, Wilda M, Kehrer-Sawatzki H, Vogel W, Fundele R, Hameister H. A high density of X-linked genes for general cognitive ability: a run-away process shaping human evolution? Trends Genet. 2001;17(12):697–701.

28. Skaletsky H, Kuroda-Kawaguchi T, Minx PJ, et al. The male-specific region of the human Y chromosome is a mosaic of discrete sequence classes. Nature. 2003;423(6942):825–837.

29. Arnold AP, Reue K, Eghbali M, et al. The importance of having two X chromosomes. Philosophical Transactions of the Royal Society B: Biological Sciences. 2016;371(1688):20150113. doi:10.1098/rstb.2015.0113

30. Cox KH, Bonthuis PJ, Rissman EF. Mouse model systems to study sex chromosome genes and behavior: relevance to humans. Front Neuroendocrinol. 2014;35(4):405–419.

31. Barker JM, Torregrossa MM, Arnold AP, Taylor JR. Dissociation of Genetic and Hormonal Influences on Sex Differences in Alcoholism-Related Behaviors. Journal of Neuroscience. 2010;30(27):9140–9144.

32. Abatan OI, Welch KB, Nemzek JA. Evaluation of saphenous venipuncture and modified tail-clip blood collection in mice. J Am Assoc Lab Anim Sci. 2008;47(3):8–15.

33. Melendez RI, Middaugh LD, Kalivas PW. Development of an alcohol deprivation and escalation effect in C57BL/6J mice. Alcohol Clin Exp Res. 2006;30(12):2017–2025.

34. Morales M, McGinnis MM, McCool BA. Chronic ethanol exposure increases voluntary home cage intake in adult male, but not female, Long–Evans rats. Pharmacol Biochem Behav. 2015;139:67–76.

35. Li J, Chen P, Han X, et al. Differences between male and female rats in alcohol drinking, negative affects and neuronal activity after acute and prolonged abstinence. Int J Physiol Pathophysiol Pharmacol. 2019;11(4):163–176.

36. Tarttelin MF, Gorski RA. Variations in food and water intake in the normal and acyclic female rat. Physiol Behav. 1971;7(6):847–852.

37. Krause EG, Curtis KS, Davis LM, Stowe JR, Contreras RJ. Estrogen influences stimulated water intake by ovariectomized female rats. Physiol Behav. 2003;79(2):267–274.

38. Santollo J, Daniels D. Control of fluid intake by estrogens in the female rat: role of the hypothalamus. FrontSystNeurosci. 2015;9:25.

39. Rodd ZA, Bell RL, Sable HJK, Murphy JM, McBride WJ. Recent advances in animal models of alcohol craving and relapse. Pharmacol Biochem Behav. 2004;79(3):439–450.

40. Vengeliene V, Bilbao A, Spanagel R. The alcohol deprivation effect model for studying relapse behavior: a comparison between rats and mice. Alcohol. 2014;48(3):313–320.

41. Füllgrabe MW, Vengeliene V, Spanagel R. Influence of age at drinking onset on the alcohol deprivation effect and stress-induced drinking in female rats. Pharmacol Biochem Behav. 2007;86(2):320–326.

42. Tambour S, Brown LL, Crabbe JC. Gender and age at drinking onset affect voluntary alcohol consumption but neither the alcohol deprivation effect nor the response to stress in mice. Alcohol Clin Exp Res. 2008;32(12):2100–2106.

43. Roberts AJ, Smith AD, Weiss F, Rivier C, Koob GF. Estrous cycle effects on operant responding for ethanol in female rats. Alcohol Clin Exp Res. 1998;22(7): 1564–1569.

44. Priddy BM, Carmack SA, Thomas LC, Vendruscolo JCM, Koob GF, Vendruscolo LF. Sex, strain, and estrous cycle influences on alcohol drinking in rats. Pharmacol Biochem Behav. 2017;152:61–67.

45. Radke AK, Sneddon EA, Monroe SC. Studying Sex Differences in Rodent Models of Addictive Behavior. Curr Protoc. 2021;1(4):e119.

46. Radke AK, Sneddon EA, Frasier RM, Hopf FW. Recent Perspectives on Sex Differences in Compulsion-Like and Binge Alcohol Drinking. Int J Mol Sci. 2021;22(7):3788.

47. Eskay RL, Ryback RS, Goldman M, Majchrowicz E. Effect of chronic ethanol administration on plasma levels of LH and the estrous cycle in the female rat. Alcohol Clin Exp Res. 1981;5(2):204–206.

48. Sanchis R, Esquifino A, Guerri C. Chronic ethanol intake modifies estrous cyclicity and alters prolactin and LH levels. Pharmacol Biochem Behav. 1985;23(2):221–224.

49. Alfonso M, Durán R, Marcó J. Ethanol-induced alterations in gonadotrophins secretion during the estrous cycle of rats. Alcohol Alcohol. 1993;28(6):667–674.

50. Burgoyne PS, Arnold AP. A primer on the use of mouse models for identifying direct sex chromosome effects that cause sex differences in non-gonadal tissues. Biol Sex Differ. 2016;7:68.

51. Arnold AP. Mouse models for evaluating sex chromosome effects that cause sex differences in non-gonadal tissues. J Neuroendocrinol. 2009;21(4):377–386.

52. Ngun TC, Ghahramani N, Sánchez FJ, Bocklandt S, Vilain E. The genetics of sex differences in brain and behavior. Front Neuroendocrinol. 2011;32(2):227–246.

53. Arnold AP. A general theory of sexual differentiation. J Neurosci Res. 2017;95(1-2):291–300.

54. Quinn JJ, Hitchcott PK, Umeda EA, Arnold AP, Taylor JR. Sex chromosome complement regulates habit formation. Nat Neurosci. 2007;10(11): 1398–1400.

55. Martini M, Irvin JW, Lee CG, Lynch WJ, Rissman EF. Sex chromosome complement influences vulnerability to cocaine in mice. Horm Behav. 2020;125:104821.

56. Carruth LL, Reisert I, Arnold AP. Sex chromosome genes directly affect brain sexual differentiation. Nat Neurosci. 2002;5(10):933–934.

